# Inter-trial variations in EEG predict the individual differences in social tasks

**DOI:** 10.1101/2021.10.29.465647

**Authors:** Haoming Zhang, Kunkun Zhang, Ziqi Zhang, Mingqi Zhao, Quanying Liu, Wenbo Luo, Haiyan Wu

## Abstract

People experience events and form an impression of others in a way that is affected by social influence every day. In the present study, we designed a series of tasks centered on social influence to investigate people’s bias in following others’ opinions and its underlying neural predictors. Our results showed social conformity and proved that social influence-induced change can be predicted by the amount of inter-trial electroencephalogram (EEG) variations when people view others’ faces. This prediction effect is robust in the alpha-band over the right frontal and left occipital electrodes for negative influence. Inter-trial EEG variations can also predict the subsequent trust difference between negatively and positively influenced faces. Our findings suggest that higher Inter-trial EEG variations in the pre-influence task may serve as a predictor of high vulnerability to social influence. The present study provides a novel approach that considers both the stability of one’s endogenous EEG and the variations in external task components to predict human social behaviors.

## Introduction

Since “beauty is in the eye of the beholder,” one’s perception or impression of faces is usually subjective (*1*). This suggests that norms exist in the social perception of faces, which can be highly similar across social contexts. However, previous research has indicated that facial judgment is modulated by various factors, such as personality (*2*), characteristics of self-resemblance characteristics (*3*), and culture (*4*). Therefore, facial judgment is subjective and comprises individual differences. Just as people do not all enjoy the same scenes or like the same characters in a movie, individuals’ thoughts and feelings may occasionally deviate from each other, reflecting variations when experiencing social stimuli. These variations in endogenous signals can impact cognitive appraisals of unfolding events and the generation of affective meaning. Consequently, individuals’ neural responses may diverge throughout a socially influenced experience, reflecting dynamically shifting alignments of thoughts and feelings. These neural dynamics may reflect the states of stable opinions or feelings that can predict people’s behavioral and neural responses when external social information is presented. How can people form a broad impression of others given a combination of these unique endogenous states and external social information?

“Social conformity” refers to people’s tendency to adjust to social influence, including others’ opinions, choices, and attitudes (*5–7*). Studies have also suggested that facial evaluation shifts due to social influence in facial judgment experiments (*8, 9*). Further, according to Schnuerch (*10*), when weaker faces are encoded, the more people conform to the majority response regarding the faces’ attractiveness—which could be attributed to the lack of reliable information on the faces. This evidence may indicate that the weaker stable norm one keeps, the more susceptibility to other’s opinions. Although evidence is mixed and still developing, prior research suggests that humans are susceptible to peer pressure (*11*). While it is evident from many psychological studies that human behavior is susceptible to the influence of others (*5, 12*), few attempts have been made to investigate whether neural state stability can predict conformity and the resulting behaviors. In light of the literature showing people’s susceptibility to social influences and the valence imbalance in impression formation demonstrated by studies in the field of social psychology (*13*), we examined whether individuals exhibit individual differences in conformity during face perception and whether such differences are predictable from their neural responses (e.g., electroencephalograms [EEGs]) in an independent face task.

Neural signatures from the frontal and occipital visual areas may predict the social influence effect on face perception. Previous studies show the role of the frontal cortex in social conflict detection and resolution (*8, 14–16*). EEG studies have also confirmed that the discrepancy between individual and majority opinions induces error or conflict detection-related components, such as frontal feedback-related negativity (FRN) (*10,17–19*). For example, research has shown that face evaluation tasks revealed robust frontal negativity when an individual’s opinion conflicted with the group’s opinion (*20*). Over the last decades, frontal alpha activities have been associated with affective motivational states, which suggests larger relative left-hemispheric activation during approach-related motivation, and larger relative right-frontal activation during avoidance-related motivation (*21–23*). In a social influence face perception task, the impression of faces can initially evoke approach or avoidance motivation, and social influence may then elicit conflict signals and aversive states, which can be indicated by frontal EEGs. Furthermore, studies have reported face-selective activation in the occipital cortex (e.g., fusiform face area [FFA]) (*24*), and a link between facial preference and EEG alpha power during passive viewing and active section tasks (*25*).

Tracking moment-to-moment variations in neural activity is emerging in neuroscience (*26–28*), and the neural variation is related to different aspects of human behavior (*29, 30*). It has been hypothesized that neural variability may reflect a greater dynamic range, such that larger variability is generally beneficial to adaptability and efficiency, which permits a greater range of responses to a greater range of stimuli (*27, 31*). Measures of brain signal variability demonstrate highly predictive results for brain function (*30, 32*). For example, such measures show greater variability during the upright compared to the inverted face condition, especially in the fusiform gyrus and anterior cerebellum (*30*). A series of studies have shown increased neural variability and lower behavioral variability in development, such that the younger group showed faster and more consistent performances and exhibited higher brain variability across tasks (*29,31,33,34*). While most of neural variations are calculated within single trials or blocks of the experiments, some studies calculated the across-trial neural variability. Studies showed that such across-trial neural variations are related to various functional roles, such as human perception (*35–38*). However, to our knowledge, no study has investigated the functional role of neural variability in social behavior variability or flexibility (*28*).

In the present study, we aimed to unveil the relation between the neural variation (i.e., human EEG) during both an independent social situation and the main tasks involving adaptive social function. Particularly, we focused on alpha-band EEG variations during a social conformity task, given that (i) the significance of alpha band power in emotion or motivation is well understood, especially among resting EEG studies; and (ii) the alpha power in different stages of life is thought to predict various human abilities (*39, 40*). Thus, it may serve as a good indicator of individual differences in brain states. Since no previous study has tried to extract alpha-band EEG in one task to predict behavioral performance or the EEG of subsequent tasks, the novelty of our work stems from the idea that the alpha-band EEG captured in an independent face perception task can predict one’s subsequent social perception and decisions.

In line with the revealed relationship between neural variation and behavior variation (*26,28, 37*), we hypothesized that the individuals who are with more variable neural activation would represent a more unstable mindset, which enables them to be more easily affected by social information from others. In another word, neural variation across trials can predict the effect of social conformity. To test our hypothesis, we first (1) proposed inter-trial EEG variation, a new method to quantify subject’s common neural variation between trials in the same task (similar to across-trial variation (*38*)); and (2) test whether inter-trial EEG variation can predict the pre-influence to post-influence behavioral change and the decision-making task (Tasks 2, 3, and 4) (see Figure 1). Our approach differs from conventional prediction studies in that we make a multiple task design (see Figure 1), and with prior assumptions about the variation characteristic of the EEG, can predict the subsequent mind-changing behaviors. This allows us to consider both the stability of one’s endogenous electrophysiological response and the variations in external task components used to predict human decision processes in a social context.

**Fig. 1.**
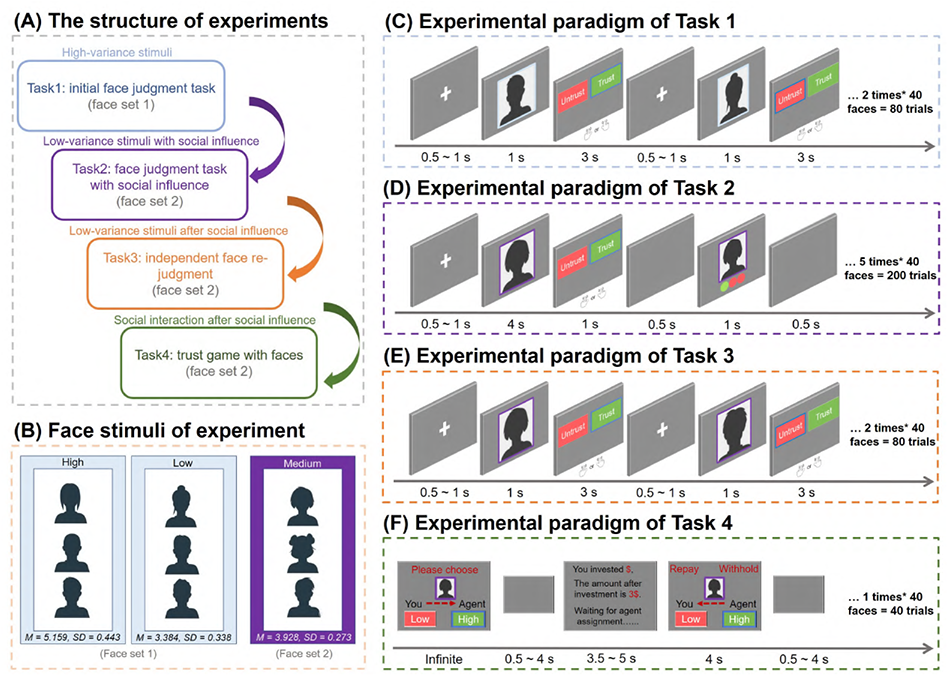
The experiment paradigm. **(A)** The structure of the four-step experimental paradigms. All participants needed to conduct four tasks successively. **(B)** Face stimuli of the experiments. Two sets of face stimuli were selected from a independent trustworthy rating experiment, controlling for some of the edge and contour features of the faces. Based on the trustworthy ratings, our experimental paradigm consisted of two types of facial stimuli, high variance in trustworthy ratings(face set 1), and low variance in trustworthy ratings(face set 2). We used face set 1 as facial stimuli in Task 1, and used face set 2 in another 3 tasks. **(C)** Experimental paradigm of Task 1. The participants need to make binary judgement of whether the presented faces is untrustworthy or trustworthy. **(D)** Experimental paradigm of Task 2. Participants were instructed to judge the neutral faces’ trustworthiness. After their responses, we will present the judgement from the other three people. **(E)** Experimental paradigm of Task 3. Participants would evaluate the faces trustworthiness again which were used in the Task 2. **(F)** Experimental paradigm of Task 4. Participants were asked to perform a trust-game experiment with each face appeared in Task 2 and Task 3.

## Results

### Behavioral results

#### Social influence altered face judgments from pre-influence to post-influence

Figure 2) presents the behavior change from pre-influence to post-influence under the four conditions. In our analysis, we considered the judgments of the first two repeated faces in Task 2 as pre-influence judgments, and the judgments in Task 3 (a judgment task without social influence) as post-influence judgments. Behavior change was defined by subtracting the average trust judgment in Task 2 (which was scored as 1 if judged as “trust,” and scored 0 if judged as “untrust”) from the average trust in Task 3 for each condition. For condition 0 (where all three peers judged as “untrust”), the average trust rate post-influence was significantly lower than the average trust rate of the pre-influence stage (*t*_31_ = 3.44*, P* = 0.0017). For condition 1 (where two peers judged as “untrust”, and one peer judged as “trust”) (*t*_31_ = 0.6897*, P* = 0.4957) and condition 2 (where one peer judged as “untrust”, and two peers judged as “trust”) (*t*_31_ = *−*0.6574*, P* = 0.5159), the average trust rate showed no significant difference between pre-influence and post-influence. For condition 3(where all three peers judged as “trust”), the average trust rate after social influence was significantly higher than the average trust rate before social influence (*t*_31_ = *−*2.483*, P* = 0.0189) (see Figure 2).

**Fig. 2.**
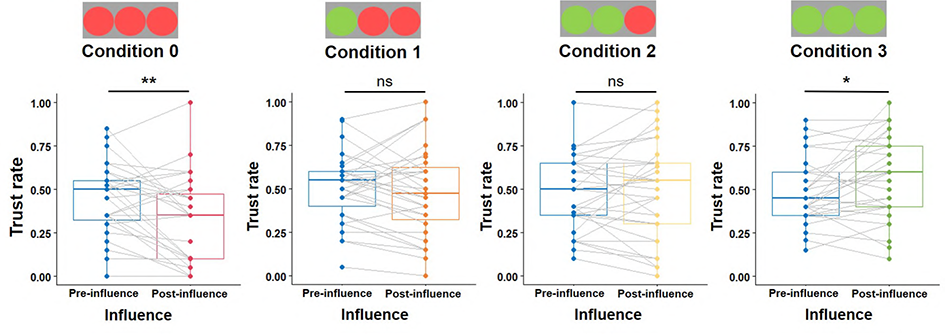
Behavioral change from pre-influence to post-influence in four conditions. Significance levels (T-test) are indicated by asterisks: \**P ≤* 0.05; \*\**P ≤* 0.01; and ns = non-significant. Condition 0: all three peers judged as “untrust”; Condition 1: two peers judged as “untrust”, one judged as “trust”; Condition 2: one judged as “untrust”, two peers judged as “trust”; Condition 3: all three peers judged as “trust”.

#### Trust rate difference between positively and negatively influenced conditions in Task 4

For the trust game in Task 4, we tested the social influence-induced behavior change, which was defined by subtracting the average trust level in conditions 0 and 1 from the average trust level in conditions 2 and 3 (Figure S1). The paired t-test indicated a significant condition difference (*t*_31_ = *−*2.431*, P* = 0.0206), and that there was a higher trust rate for positive socially influenced faces (conditions 2 and 3) than negatively socially influenced faces (condition 0 and 1).

### EEG results

#### EEG variations in Task 1

In Task 1, we first used the EEG signals from all frequency bands. According to the theory on the link between frontal alpha-band power and motivation system (i.e., left and right frontal alpha power reflects approach and withdrawal motivation respectively) (*41*), we believe that trust and untrust judgments are closely related to motivation directions. Thus, we used a bandpass filter to extract the time-domain EEG epoch signals of alpha bands (8–13 Hz).

To obtain subject-level measures of temporal variation, we first segmented the EEG epoch time series into m windows, each with a length of 20 ms. Within each time window, the inter-trial EEG variation was obtained by 1 subtracting the average value of the lower triangular correlation matrix, which was obtained by calculating the Pearson correlations between any two trials in all trials. In Figure 3, we present the correlation matrix of the subjects with the highest variation and lowest variation (electrodes AF4 and P7). We show the correlation matrix of all the subjects(electrodes AF4 and P7) in Figure S3 and Figure S4, respectively.

**Fig. 3.**
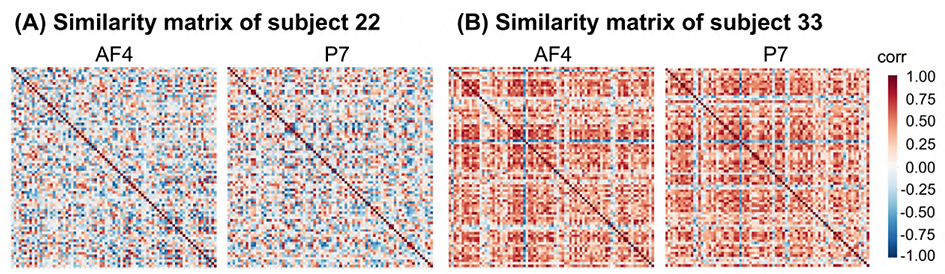
Some representative similarity matrices calculated using the EEG in Task 1. The similarity matrix was obtained by calculating the Pearson correlation of all trials in pairs, and was averaged between 0–300 ms.. **(A)** The similarity matrix of the subjects with highest variations (0-300 ms). The similarity matrix was calculated using electrode AF4 (left), and P7 (right). **(B)** The similarity matrix of the subjects with lowest variations (0–300 ms). The similarity matrix was calculated using electrode AF4 (left), and P7(right).

As shown in Figure 4, we observed a general peak variation during 150-200 ms over the right frontal electrode (AF4) and left occipital electrode (P7). Similar patterns can be observed in other electrodes in Figure S9. There was no significant difference in EEG variation between the trust and untrust conditions for electrodes AF4 or P7, so we obtained the mean variation for each participant across the two conditions.

**Fig. 4.**
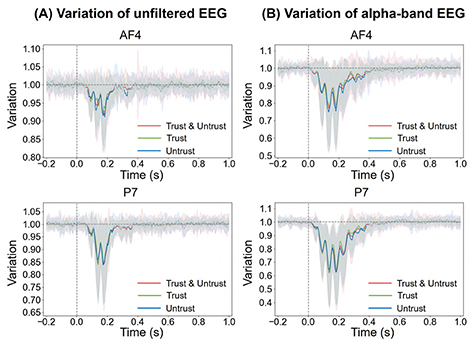
Some representative epochs of the EEG temporal variation, from electrode AF4 (upper) and electrode P7 (bottom). **(A)** Variation was calculated using unfiltered EEG. **(B)** Variation was calculated using alpha band EEG. The red, green, and blue lines are the variation calculated by all events (Trust & Untrust), only the Trust event and only the Untrust event, respectively. The shaded areas represent the standard deviation. Dashed lines indicate that the variation is not significantly different from the baseline, and solid lines indicate that the variation is significantly different from the baseline.

#### EEG variations in Task 1 can predict behavioral changes from pre-influence to post-influence

After quantifying the Inter trial EEG variation in Task 1, we sought to evaluate whether the EEG variation pattern within one independent face set could predict the following social influence effect in the face perception of another face set. We applied the correlation analysis, using the EEG variations of each electrode in Task 1 to predict the behavioral change from pre-influence to post-influence. We also conducted statistical significance tests along with the correlation analysis, and only saved significant(*P <* 0.05) correlation results. To visualize our results, we plotted the electrodes that were significantly correlated with behavioral changes on a topographic map (see Figure 5A). We found that EEG variation at a large number of electrodes was significantly negatively correlated with behavioral change over a longer period of time in conditions 0 and 1. The negative correlation indicates that the higher the EEG variations in Task 1, the more susceptible they are to the negative social influential effect in Task 2. For condition 0 (see Figure 5A, upper), the electrodes in the right frontal and left occipital lobe show a significantly negative correlation from 25 ms to 160 ms. For condition 1 (see Figure 5A, bottom), the electrodes in the right frontal and left occipital area showed a significantly negative correlation from 35 ms to 300 ms.

**Fig. 5.**
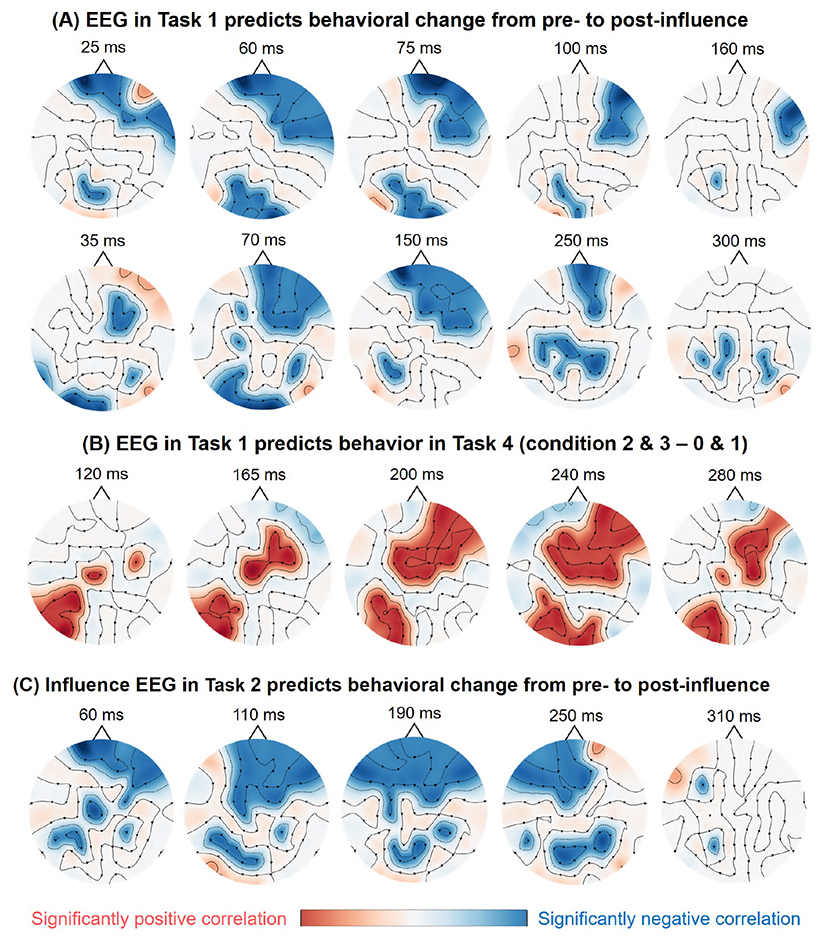
The topographic maps shows that the EEG variation in some brain regions can be significantly correlated with behavior. The red and blue areas on the topographic map represent that the EEG variation in these areas are significantly positively and negatively correlated with behavior, respectively. **(A)** EEG variation in Task 1 can predict the behavioral change from pre-influence to post-influence in condition 0 (upper) and condition 1 (bottom). **(B)** EEG variation in Task 1 can predict the difference between conditions 2 and 3 as well as conditions 0 and 1 in Task 4. **(C)** EEG variation under social influence in Task 2 can predict the behavioral change from pre-influence to post-influence in condition 1.

#### EEG variations in Task 1 can predict trust rate in Task 4

Next, we tested whether the EEG variations in Task 1 could predict the subsequent social influence effect in Task 3. Similar to the previous analysis, we calculated the correlation of EEG variation and behavior difference at each electrode and conducted statistical significance tests. We found that the electrodes in the right frontal, right parietal, and left occipital lobe showed a significantly positive correlation from 120 ms to 280 ms (see Figure 5B). This positive correlation means that the higher the EEG variations in Task 1, the higher the trust rate difference between positively and negatively influenced conditions in Task 4.

#### EEGs under the influence of Task 2 can predict behavioral changes from the pre-influence to post-influence stage

Since the individual EEG revealed that variations can predict behavioral change from pre-influence to post-influence, we asked that whether the EEG variations in others’ evaluations would exhibit a predictive effect on subsequent behavioral change? Using the same analysis method, we found that the electrodes over the frontal lobe showed a significantly negative correlation with behavioral changes from 60 ms to 310 ms (see Figure 5C). This negative correlation means that the higher the EEG variations under the influence of Task 2, the more susceptible they are to negative social influence effects in Task 2. Additionally, the correlated map showed a temporal tendency to move from the right frontal lobe to the left frontal lobe. At approximately 110 ms, the right prefrontal lobe showed a significantly negative correlation with behavior; and at 190 ms, the whole prefrontal lobe showed a significantly negative correlation; and at 250 ms, the left prefrontal lobe showed a significantly negative correlation.

#### EEG variations in Task 2 and 3

After evaluating the temporal EEG variation in Task 1, we also evaluated the temporal EEG variation when the participants judged the faces in Task 2 and 3. We wondered whether the EEG variations in Task 2 and 3 would have the same temporal pattern as the EEG variations in Task 1. The method used to calculate the variation of Task 2 and 3 was similar to that of Task 1. To explore whether the EEG variation would gradually change as the repetition of influence increases, we calculated EEG variations for each condition and each influence step independently.

In previous results, we have found that the region of the right frontal and left occipital lobe was significantly correlated with behavioral changes. To focus on the region of the right frontal and left occipital lobe, we divided the 63 electrodes into six regions of interest (see Figure 6A) (*42*). We considered the region 2 as the region of right frontal area, and the region 5 as the region of left occipital area. The variations in each region were calculated by averaging the variations across all electrodes within the given region.

**Fig. 6.**
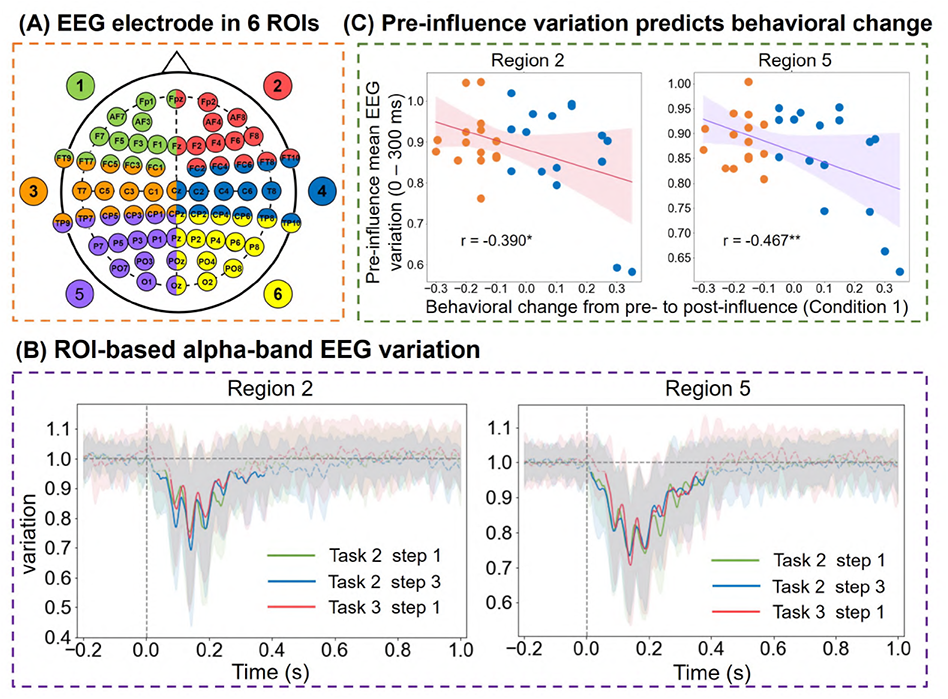
EEG variations in pre-influence judgement state can predict behavioral change. **(A)** EEG electrode in 6 regions of interest (ROIs). The 63 electrodes were grouped into six regions. We included the electrodes in the border of two regions in both regions. **(B)** Some representative epochs of the EEG temporal variation in Task 2 and Task 3, region 2 (left) and region 5 (right). The green, blue, and red lines are the EEG variation calculated by the first time the face was presented in Task 2, the third time the face was presented in Task 2, and first time the face was presented in Task 3, respectively. The shaded areas represent the standard deviation. Dashed lines indicate that the variation is not significantly different from the baseline, and solid lines indicate that the variation is significantly different from the baseline. **(C)** The mean EEG variations in Task 2 step 1 - step 3 can predict the behavior change from pre-influence to post-influence. The left side and right side are the mean EEG variations of region 2 and region 5, respectively. The yellow dots are the subjects who were most negatively influenced. The blue dots are the subjects with little or no negative impact. Significance levels (T-test) are indicated by asterisks: \**P ≤* 0.05; \*\**P ≤* 0.01.

We show the correlation matrix of all the subjects in Task 2 (see Figure S5 for region 2, Figure S6 for region 5) and Task 3 (see Figure S7 for region 2, Figure S8 for region 5). Additionally, Figure 6B presents the EEG variation when the face set was shown for the first time in Task 2, the third time in Task 2, and the first time in Task 3 as an example. The EEG variations in Task 2 and Task 3 showed a similar pattern to the EEG variations of Task 1. We performed a cluster-level statistical permutation test on the temporal EEG variation of different influence steps and found that there was no significant EEG variation difference as the number of influence steps increased.

#### EEG variations during the pre-influence period in Task 2 can predict behavioral changes from pre-influence to post-influence

We sought to evaluate whether the EEG variation under the pre-influence face judgment period could predict the subsequent social influence effect. Before we applied the correlation analysis and significance test, we averaged the temporal variations from 0 s to 300 ms to obtain the pre-influence mean variations. Next, we used the pre-influence mean variation of each region to predict the behavioral change from pre-influence to post-influence. For condition 0, we found that the mean variation in region 5 was significantly negatively correlated with behavioral change (*r* = *−*0.415*, P* = 0.02). For condition 1 (see Figure 6C), we found that the mean variation in regions 2 and 5 was significantly negatively correlated with behavioral changes (Region 2: *r* = *−*0.390*, P* = 0.03; Region 5: *r* = *−*0.467*, P* = 0.008). For conditions 2 and 3, there was no significant correlation between the mean variation and behavioral change.

## Discussion

Although the understanding of the neurobiological processes associated with social conformity has increased over the years, few studies have attempted to examine whether there is individual differences in neural signals can predict people’s sensitivity to social conflict and social conformity. Our study first examined whether the effect of social influence in the face perception task could be predicted by the EEG in a higher variance stimuli task. Consistent with our hypothesis, we find predictive effect for the social conformity of individuals with their inter-trial EEG variations.

### Why can EEG variations during high-variance stimuli can predict the social influence effect?

The human brain processes complex social information in both stable and flexible ways (*43, 44*), integrating speed and efficiency. Even when facing with high variations in external stimuli, a the stable neural system directly processes social information that allows us to generate and maintain a sense of social perception. The fast social brain response is likely higher similar across individuals who share the same social experience (*45, 46*). However, given the same social information, individuals with different social experiences may have unique interpretations, feelings, and decisions-often leading to differed social cognition across individuals (*47, 48*). How can people establish a flexible social functional system that adapts to dynamic social environments, and how can such neural fluctuations in social contexts across individuals independently predict susceptibility to social influence or adjustment under social conflict? Our evidence shows that the stability or variations in specific experiences may be unique to each individual and predict their responses in a social context.

Although it is of considerable importance, how human social behavior can be predicted from task-dependent EEG variations, is still unclear. Primarily, no existing study links pre-task EEG variations to individual differences in subsequent social behavior. The presently used EEG neural distinction between trustworthy and untrustworthy faces may reflect a social perception organizing principle of the brain, which may originate from past social experience or a stable social perception system. Since the decoding accuracy using pattern classification is not significantly higher than the baseline in Task 1 (see Figure S2), the EEG signal appears to be more variable across individuals than across conditions. The strong correlation between the EEG variation of Task 1 and the EEG variation of pre-influence stage of Task 2 further proves this point (see Figure S11). Moreover, the initial distinction between the two types of faces can predict the subsequent rating changes under social influence. Unlike previous EEG alpha power extracted from resting EEG, or or the prediction of single-task performance, we provide the first evidence that task-dependent drift for high-variance stimuli can predict social impression changes. That is, the degree to which people are vulnerable to the “disturbance” of peer pressure is predictable from a combination of individual EEG variations.

### Prediction effect lasts: EEG in Task 1 can predict decisions after social influence

Studying the stable prediction of the effect is generally involves two ways. One to test the prediction effect across different samples (*49, 50*), and the other way is to test the prediction effect across different time points or tasks (*51, 52*). Neural variability exhibits stability, both across different task conditions, as well as across trials with the same task condition. An recent view proposed shared variability is associated with learning, that “behavioral variability is driven by neural variability” (*53*). Unlike previous studies that captured EEG features from the resting state, we defined the task-evoked EEG variations to predict subsequent social behaviors, across tasks.

In our findings, the “behavioral variability” induced by the social influence persists to the trust task(higher trust rate for positively influenced faces), even the social influence stimuli is removed in the task. Further, Ithe prediction effect manifested not only on socially influenced changes but also from pre-task EEG variations to trust decisions after the influence task. It showed stable predictability over longer periods, supporting the relative stability of variations. The task evoked behavioral change effect, both the rating change and decision difference to the influenced faces, supports the relative stability of behavioral variability. These prediction effects further support the stable link between behavioral variations and neural variations. These prediction effects of alpha oscillation “bridge the gap” between task-dependent EEGs and task-induced social behaviors, and provide novel insights into how task-dependent EEGs predict behavior and cognition. They may likewise contribute to the prediction of different aspects of social cognition, such as stereotypes, discrimination, and in-group/out-group effects.

### Specificity of prediction effect: EEG over right frontal and occipital can predict following social behaviors

EEG oscillations have also been linked to the prediction of human behavior. The alpha frequency between 8 and 12 Hz was the dominant frequency of the human EEG. In our findings, the prediction effect primarily occurred over the right frontal and occipital areas. Such consistent prediction effect specificity may be interpreted from two accounts: 1) the social task specificity, and 2) the interplay of frontal and occipital areas suggests task-general flexibility and stability in prediction (*54*).

For the first account, frontal activity is related to reinforcement learning(RL), errors, and conflict processing (see a review by (*55, 56*)). For instance, in a probabilistic reinforcement learning task, a single-trial analysis revealed that the frontal theta band correlated with reward prediction error (both negative and positive errors) (*57*). It is also suggested that different pre-frontal regions are activated by different types of prediction errors, and the frontal alpha in our results may also reflect this social influence task-dependent activity that can predict the behavioral adjustment after exposure to social information. Since the main task focuses on faces, the prediction effect over the occipital area fits both our hypothesis as well as previous research indicating an association between alpha and the visual representation of the face (*58*).

On the other hand, since we observed the prediction effect across tasks, we tend to support various task-nonspecific processes. The the prediction alpha activity may also reflect task-general mind wandering (*54, 59*), or a general integration of flexibility and stability in prediction, as previous research has shown the frontostriatal contribution to the such interplay (*60*). It is a challenge to give a conclusion on this point given that our tasks are social influence-related processes, resulting in difficulties in generalization to other domains. Further prediction studies on different social or cognitive tasks should be conducted to validate these accounts.

### Prediction effect difference across conditions: Higher behavioral changes and predictability for negative social influence conditions

In the face judgment task, the participants received both positive and negative social influences from their peers. We consistently found a more pronounced effect among negative influence conditions, both on behavioral change and EEG prediction. We believe that a prominent explanation for these effects is the negativity bias in social information processing. The behavioral change from Task 2 to Task 3 (Figure 2) indicated a more pronounced effect and impression updating in the negative influence condition. This supports the presence of a negativity bias that has a greater impact on negative evaluations, compared to available positive evaluations. Positive-negative asymmetry has already been demonstrated in the context of evaluation, attitude, and impression formation (*61–64*). In an early study on using adjectives to describe another person, participants placed more weight on negative descriptions of others than on positive ones (*65*). Furthermore, our findings support the theory proposed of Rozin and Royzman (*66*), who stated regarding negativity dominance, “if we then find that losing $100 is as bad as winning $150 is good, and that losing $100 and winning $150 is negative, then we have negativity dominance.” Interestingly, participants showed higher predictability for the negative influence condition—indicating that people weigh negative social influence more in the social context. This range to shift to a more negative impression or negativity weight can be better captured or predicted by EEG variations. This further indicates that salient social information processing would be better predicted with task-dependent EEGs.

Overall, the present study provides further evidence that negative information from others has a stronger impact than positive information. It likewise provides the temporal profile of social conflict processing, as revealed by alpha-band EEG and oscillation patterns.

#### EEGs during social influence predict rating changes

The reinforcement learning theory of social conformity holds that the differences between outcomes and expectations (i.e., prediction errors) play a crucial role in guiding individual adaptive behavior. Outcomes better than expected reinforce behaviors, while outcomes worse than expected signal a need for adjustment (*67, 68*). Accordingly, the mismatch between one’s own evaluation and group ratings may be detected as a “norm prediction error,” indicating a need to correct for deviance from norms (*8, 14, 69, 70*). Following our previous EEG results (*71*), such error detection signals can be indexed by using the frontal negativity. The present findings further confirm the activation of frontal electrodes in awareness of social conflict or social deviation, which may subsequently predict behavioral change (Figure 5C). Alpha and theta band activity has also been well-documented as an oscillation signal associated with awareness and cognitive conflict. Frontal area alpha band activity indicates activity in the anterior cingulate cortex (ACC) (*72*) and is expected to predict behavioral adjustment. Although we did not examine the alpha band power difference between the negative and positive conditions, we provided further evidence of the predictability of negative influenced conditions. We found that alpha band variation in the Task 1, and pre-influence stage of Task 2 correlated with the behavioral changes. This indicated that the variation of alpha band neural activity may be associated with the flexibility of social adaptation, which predicts subsequent impression change. This is also evidence of the tuning of one’s internal evaluation system toward external social inputs, as the perceived social conflict could predict such integration of a stable personal state and various external social stimuli.

In conclusion, our results provide novel insights into the role of EEG variations in the prediction of social perception and decisions. The present study shows that the prediction effect exists and is more pronounced for more salient information or brain regions, complementing previous research that has implicated EEG indicators in behavior prediction. Our approach is of significance for various social behavior predictions from EEG variation and detects social stability and social flexibility in humans.

## Materials and Methods

### Participants

Thirty-four participants took part in the experiment. Data from three participants were excluded from the analysis due to data loss or EEG artefacts. The remaining 31 participants (17 females, 14 males), aged between 19 and 25 years old (*M* = 21.45, *SD* = 1.959), were included in the analysis. There was no difference in the age range for female (*M* = 21, *SD* = 1.936) and male participants (*M* = 21.818, *SD* = 1.991). All participants were right-handed, had normal or corrected-to-normal vision and reported no history of neurological or psychiatric disease. All participants provided written informed consent and were paid 100 Chinese yuan for their participation in the study.

### Face stimuli

Facial stimuli were obtained from the CAS-PEAL face database (*73*). First, a total of 160 faces were rated on a seven-point scale (from “1” very untrustworthy to “7” very trustworthy) by 30 participants recruited independently from the formal study. Based on these ratings, we selected two sets of faces that 40 faces for Task 1 (face set 1), and another 40 faces for Task 2, Task 3, and Task 4 (face set 2). Specifically, in the face set 1, we included 20 trustworthy faces (*M* = 5.159, *SD* = 0.443) and 20 untrustworthy faces (*M* = 3.384, *SD* = 0.338) for Task 1. The ratings of trustworthy face are significantly higher than the untrustworthy faces (*p <* 0.01). For face set 2, we selected 40 middle-level trustworthy faces (ratings ranging from 3 to 5). Each face was presented five times in the Task 2, twice in the Task 3, once in the Task 4. The average ratings of the faces in face set 2 were neutral (*M* = 3.928*, SD* = 0273). Notably, the variance and range of trustworthiness in face set 1 are larger than those in face set 2.

### Experiment design and procedure

The participants sat in a comfortable chair in a dimly lit room facing a computer monitor at a distance of 60 cm. After filling out the informed consent form, the electrode cap was mounted, and the experiment instructions were clearly shown to the participants. They were asked to minimize blinking and maintain visual fixation on a small cross in the center of the screen during the experiments. Each participant performed approximately 10 practice trials before the formal experiments. The full study consisted of four sequential experiments.

#### Task 1: Initial face rating task

Task 1, the initial face rating task, consisted of 40 trials (Figure 1C). Face set 1, which comprised two levels of trustworthiness, was presented in a randomized order in each trial. The participants had to make a binary decision on whether the presented face was untrustworthy (press “F” with the left index finger) or trustworthy (press “J” with the right index finger). Each face is presented for twice in Task 1. Behavioral data (*e.g.,* decision time and decisions) and EEG signals were recorded.

#### Task 2: Social influence task

Task 2 comprised 200 trials (Figure 1D) and consisted of presenting facial set 2 to the participants. Each trial consisted of two rounds of binary decision-making. First, the participants made a binary decision on whether the presented face was untrustworthy (press “F”) or trustworthy (press “J”); this was the pre-influence round. The trustworthy levels given by three peers were then presented for 1 s. In the post-influence round, the participant had to make a second decision on the trustworthy level of the face. To illustrate the judgment made by the three other people, three color-coded circles were displayed simultaneously below the face judged by the participant. The circle was displayed in green or red, in order to reflect whether the same face was identified as trustworthy (color = green) or untrustworthy (color = red) by other people. . Thus, there were four types of color-coded circle combinations: three red circles, which means all three people rated the face as an untrustworthy face; two red circles and one green circle, which means two people rated it as an untrustworthy face and one rated it as a trustworthy face; one red circle and two green circles, which means one person rated it as an untrustworthy face and two rated it as a trustworthy face; and three green circles, which means all three people rated it as a trustworthy face. In this manipulation, there were four conditions (10 faces for each condition), and the participants judged a series of 40 faces five times. In each trial, the participant viewed the screen showing a photograph of a neutral face, and the responses had to be made by pressing one of two keys(untrustworthy face: “F”, trustworthy face: “J”) with the left and right index finger. Upon responding, the ratings of their peers were shown for 3 s, and the next trial commenced. Note that for each face, the other’s rating is kept the same (i.e., each face displays the same three color-coded circles).

#### Task 3: Re-judgements of face set 2

To test the left-over effects of social influence in Task 2, each face was re-judged twice in Task 3 (Figure 1E). No social influence was presented in this session.

#### Task 4: Trust game with face set 2

To test whether the social influence in Task 2 would change the general trustworthiness of the face, rather than only has a rigid memory of the trustworthy level of each face, we design Task 4, a multi-round trust game. Task 4 consists of 40 trials (Figure 1F), each face in face set 2 was presented once and participants have to choose to invest a certain amount of money to themselves and the presented agent. In each trial, the participant first needs to make investment choices for the agent press “F” represents low investment, and press “J” key represents high Investment. Then, a message will be displayed in the middle of the screen: *“You invested 1$ to the agent, and the amount after investment is 3$. Please wait for the agent to allocate the amount…”*. After that, the left side of the screen shows the amount the agent repay to the subject, and the right side shows the amount the agent withhold for himself. At the end of each trial, the participants need to rate their happiness of each faces’ repayment offer.

### EEG data acquisition and preprocessing

The EEG was continuously recorded from 63 electrodes with BP amplified using the SynAmps system. A common average reference and a forehead ground electrode were used (FCz). Vertical electro-oculographic (EOG) activity was recorded with additional electrodes located below the right eye. For all electrodes the impedance was kept less than 5 kΩ. Electrical activity was amplified with a bandpass of 0.01–100 Hz and a sampling rate of 1000 Hz. In offline analysis, the data was preprocessed using MATLAB. The data was band-pass filtered between 0.5 to 30Hz, notched at powerline frequency. Artifacts related to eye and muscle activity were detected and removed by independent component analysis (ICA). Then the data were epoched into single sweep recordings from 700 ms before to 1500 ms after stimulus onset. Moreover each epoch was baseline corrected using the signal recorded during 200 ms that preceded the onset of the stimulus. All epochs with ocular artifacts greater than 30 mV were automatically rejected and in addition were visually scanned to find further artifacts. In the last step, we used bandpass filter to extract time domain EEG epochs signal of alpha bands (8-13 Hz) as the previous theoretic on link between frontal alpha band power and motivation system (*41*).

### Calculate EEG variations

To obtain subject-level measures of inter-trial EEG temporal variation, We used a 20 ms time window to segment alpha band EEG epoch time series into *m* windows with stride equal to 1 ms (see Figure 7). We also try the time window with different lengths, for example, 10 ms and 50 ms. To balance the time resolution and the smoothness of temporal variations, we chose the 20 ms length time window. (see Figure S10) The time window starts 200 ms before the stimulus and ends 1000 ms after the stimulus. Within each time window, we calculated the similarity matrix (see Figure S3) of all trials by Pearson correlation, and averaged the lower triangle of the matrix to obtain the average EEG similarity. Next, average EEG variation is obtained by 1 subtract the average EEG similarity. (see Eq. 1)

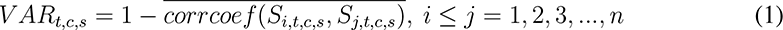

**Fig. 7.**
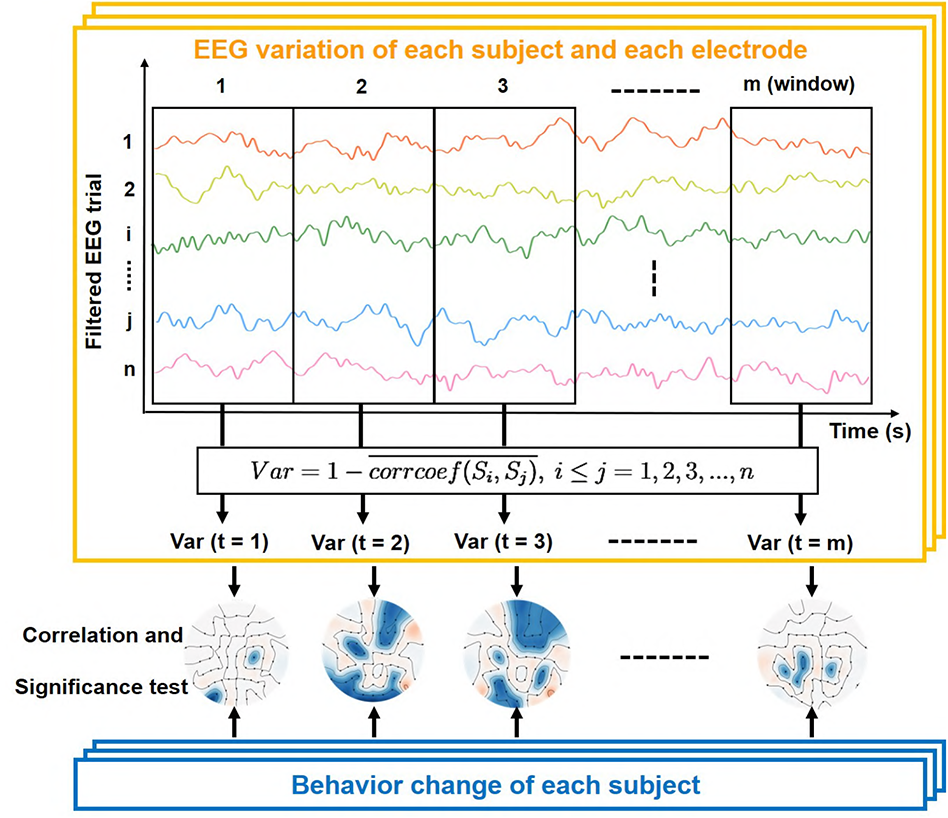
The pipeline for calculating EEG temporal variation and correlating with behavioral data. The yellow box contains the definition of temporal variation. The time series are EEG signals for all trials of each subject and each electrode. We used a time window to segment EEG epoch time series. Within each time window, we calculated the similarity matrix of all trials by Pearson correlation, and averaged the lower triangle of the matrix to obtain the average EEG similarity. Next, the average EEG variation was obtained by subtracting one from the average EEG similarity. We then calculated the Pearson correlation between EEG variation and behavioral changes of all subjects. We performed statistical significant tests to the resulting correlations, and retained the correlation results with *P <* 0.05.

Where *V AR_t,c,s_* denotes the EEG variation in time window *t*, electrode channel *c*, and subject *s*. *S_i,t,c,s_* as the EEG signal of *i^th^* trial and window *t*, electrode channel *c*, and subject *s*. *S_j,t,c,s_* as the EEG signal of *j^th^* trial and window *t*, electrode channel *c*, and subject *s*. *n* as the trial number in total. We have adopted this same measure to address the EEG temporal variation of each subject and electrode.

For the EEG variations when participants judge the faces in Task 2 and Task 3, we further divided the 63 electrodes into six regions of interest (see Figure 6A). We allocated the electrodes to six brain regions to pay more attention to the right frontal lobe (region 2) and left occipital lobe (region 5). To obtain more stable spatial patterns, we included the electrodes in the border of two regions in both regions, adapted from (*42*). The variations in each region were calculated by average variations across all electrodes in this region.

In order to calculate whether the amplitude of each EEG variation is significant relative to baseline (variation = 1), we use one-sample permutation cluster test with a p-value threshold equal to 0.05, and the number of permutations equal to 10000. To calculate whether there is significance between the magnitudes of two EEG variations, we use permutation cluster test with a p-value threshold equal to 0.05, and the number of permutations equal to 10000.

### Predictive models building

The main part of our analysis is using the EEG variations to predict the behavioral change(see Figure 7). We calculated the Pearson correlation between EEG variation and behavioral changes of all subjects. We also performed statistical significance tests along with correlation analysis, and retained the correlation results with *P <* 0.05.

For EEG variations in Task 1 and EEG variations under the influence stage of Task 2, we used a 50 ms long window with a step size of 1 ms to segment the time series of EEG variations to obtain a more stable temporal correlation pattern. For each time window, we average all points in the time window and use the average value to calculate the correlation with behavioral data. Finally, we get the correlation and p-value between the variation of each electrode at each time point and the behavioral data. To overcome the multiple comparison problem, we only focus on the significant correlation result over large areas: we only judge the significant correlation result as valid if at least three adjacent electrodes are significant simultaneously.

For EEG variations of pre-influence stage in Task 2, we select the temporal EEG variations within 0 to 300 ms for each region. We choose the EEG variations within 0 to 300 ms for two reasons. The first reason is according to the EEG variation pattern of Task 2 (see Figure 6B), the amplitude of variation significantly lower than baseline in this period of time. The second reason is in the results shown in Figure 5, the time period during which EEG variation can be used to predict behavior change is also around 0 - 300 ms. Next, we average all points in 0 - 300 ms to a average value and use the average value to calculate the correlation with behavioral data. Finally, we get the correlations and p-values between the variation of each region and the behavioral data.

The corresponding codes for calculating EEG variation are available online at https://github.com/andlab-um/Trust-untrust-face-judge.

## Acknowledgements

The authors thank Dr. Yanyan Qi for helpful comments.

## Author contributions

HW conceived the research. KZ and ZZ conducted the experiment. HZ and HW designed the analyses. HZ, MZ and HW conducted the analyses. All authors wrote the manuscript.

## Data and materials availability

Additional data and materials are available online.

## Supplementary Materials

### 1 Trust rate difference between positively versus negatively influenced conditions in Experiment 4

As shown in Figure S1, we plotted the trust rate for negatively influenced conditions (condition 0 & 1) and positively influenced conditions (condition 2 & 3). It indicated that the trust rate is higher for positively influenced conditions(M = 0.531, SD = 0.251) than for negatively influenced conditions (M = 0.604, SD = 0.244)

### 2 MVPA for EEG in Experiment 1

In Figure S2, we presented the results of MVPA, while they were not significant at the correction level (*p <* 0.05). We trained and tested series of linear estimators on brief (20 ms) consecutive windows all along the time course of the alpha band EEGs.

### 3 EEG variation calculated by the time window of different length

In our result, we calculate the EEG variations by 20 ms time window. We did not choose the 20ms window arbitrarily, we also try the time window with different lengths, for example, 10 ms and 50 ms. (see Figure S10) The temporal variations calculated by 50 ms time window are smoother than the temporal variations calculated by 20 ms time window, but the poorer temporal resolution (see Figure S10 A). The temporal variations calculated by the 10 ms time window have better temporal resolution but more rough (see Figure S10 B). To balance the time resolution and the smoothness of temporal variations, we chose the 20 ms length time window.

### 4 EEG variation calculated by the other frequency band

We have repeated the same analysis method to the EEG signal filtered by other frequency bands, for example, delta (0-4 Hz), theta (4-8 Hz), and beta (13-30 Hz), and found no meaningful result.

### 5 Compare the EEG variations of Experiment 1 with the EEG variations of Experiment 2 and Experiment 3

Since both the EEG variation of Experiment 1 and Experiment 2 during pre-influence period can predict behavior changes, we sought to evaluate whether the EEG variations in experiment 1 can correlate with the EEG variations under the pre-influence judgement of experiment 2. To compare with EEG variation of experiment 2, we divided the 63 electrodes of experiment 1 into six regions. As shown in Figure S11, the result indicated significantly correlation in all 6-regions (Region 1: *r* = 0.769*, P* = 4.44 *∗* 10*^−^*^7^; Region 2: *r* = 0.701*, P* = 1.12 *∗* 10*^−^*^5^; Region 3: *r* = 0.869*, P* = 2.39 *∗* 10*^−^*^10^; Region 4: *r* = 0.803*, P* = 5.35 *∗* 10*^−^*^8^; Region 5: *r* = 0.832*, P* = 6.58 *∗* 10*^−^*^9^; Region 6: *r* = 0.852*, P* = 1.22 *∗* 10*^−^*^9^)

---

**Figure S1:**
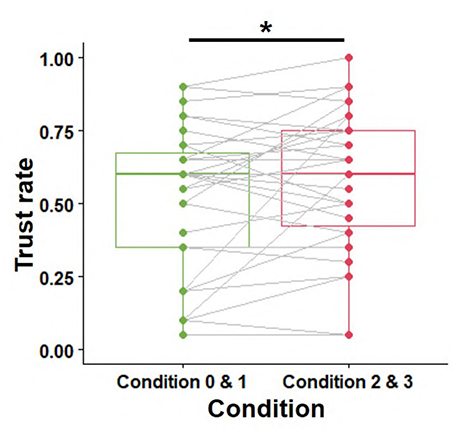
Behavioral result of Experiment 4. As described in the text, from left to right is the mean trust rate for faces of condition 0 & 1, the trust rate for faces of condition 2 & 3. Significance levels (Paired-Samples T-test) are indicated by asterisks: *p *≤* 0.05; **p *≤* 0.01; and ns: non-significant.

**Figure S2:**
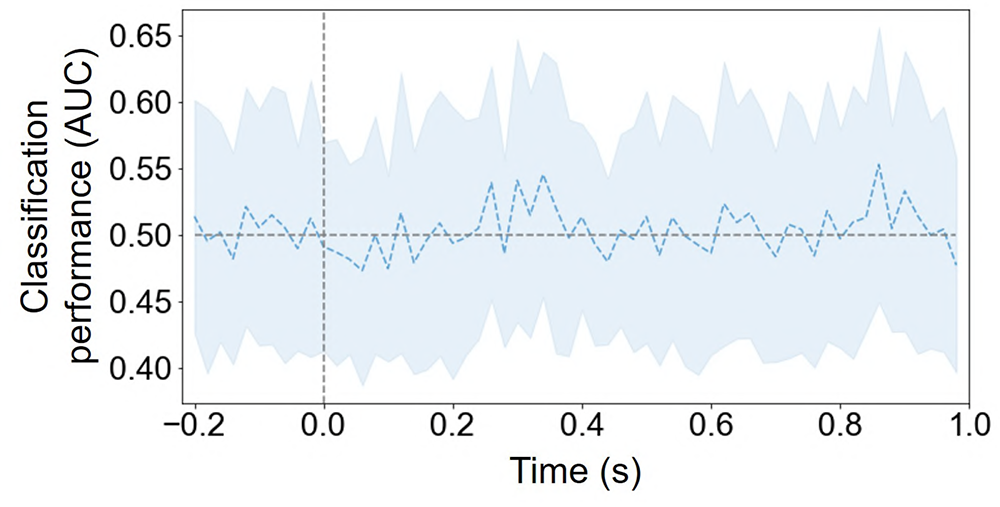
Classification performances of estimators trained on single time windows (20 ms) along the alpha band EEG. The shaded areas represent the standard deviation. Dashed lines indicate that the performance is not significantly different from the change level (AUC = 0.5)

**Figure S3:**
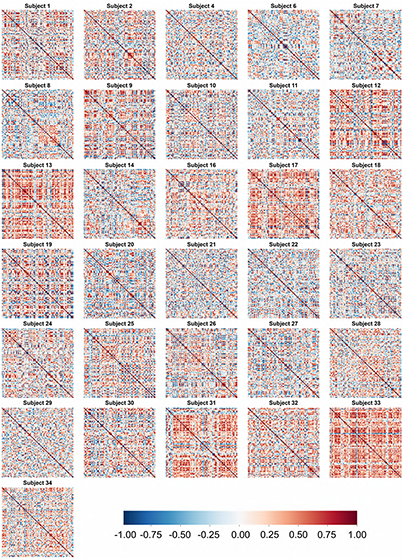
The similarity matrix of all subjects with electrode AF4 in Experiment 1.

**Figure S4:**
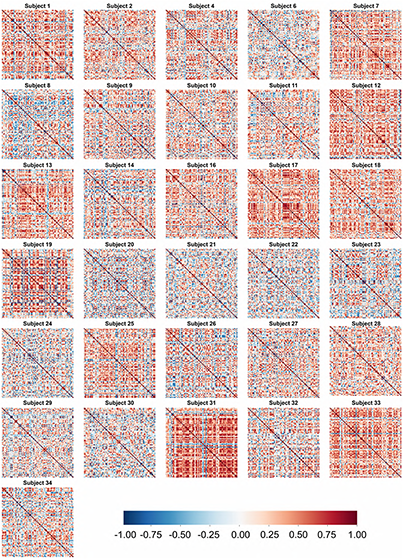
The similarity matrix of all subjects with electrode P7 in Experiment 1.

**Figure S5:**
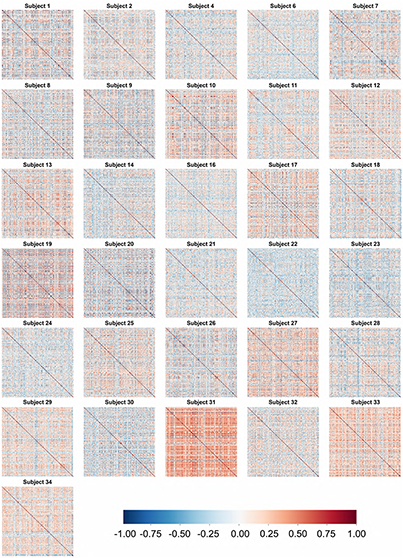
The similarity matrix of all subjects with region 2 in Experiment 2.

**Figure S6:**
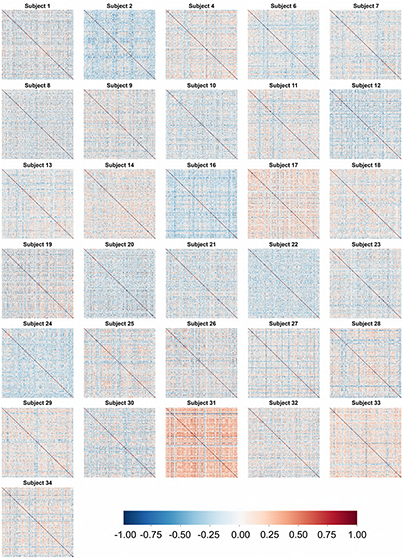
The similarity matrix of all subjects with region 5 in Experiment 2.

**Figure S7:**
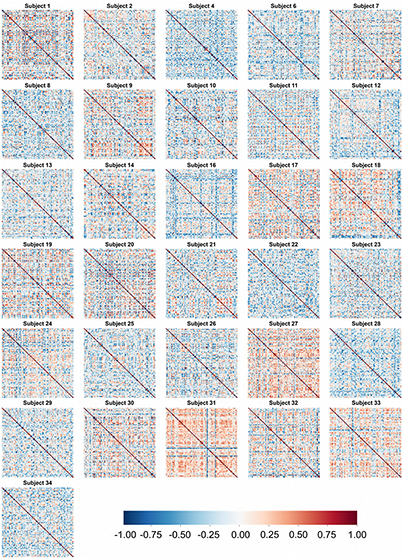
The similarity matrix of all subjects with region 2 in Experiment 3.

**Figure S8:**
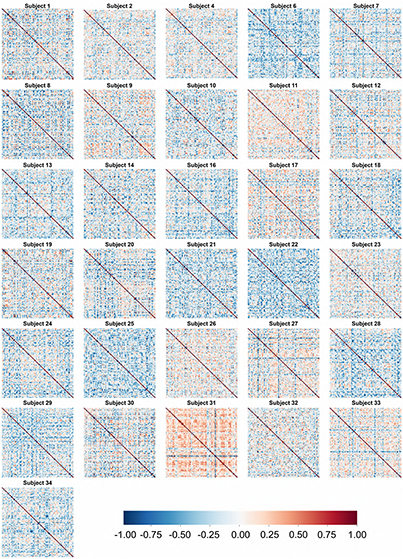
The similarity matrix of all subjects with region 5 in Experiment 3.

**Figure S9:**
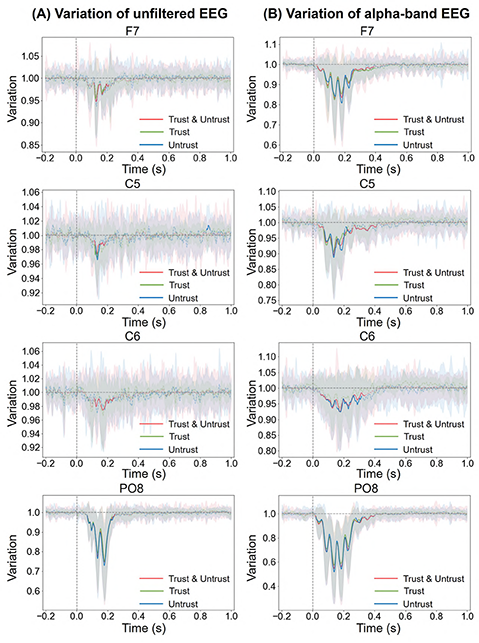
Some representative epochs of the EEG temporal variations. The electrode name from upper to bottom: F7, C5, C6, PO8. (A) Variations calculated by unfiltered EEG. (B) Variations calculated by alpha band EEG. The red, green and blue line are the variability calculated by all events (Trust & Untrust), only Trust event and only Untrust event, respectively. The shaded areas represent the standard deviation. Dashed lines indicate that the variability is not significantly different from the baseline, and solid lines indicate that the variability is significantly different from the baseline.

**Figure S10:**
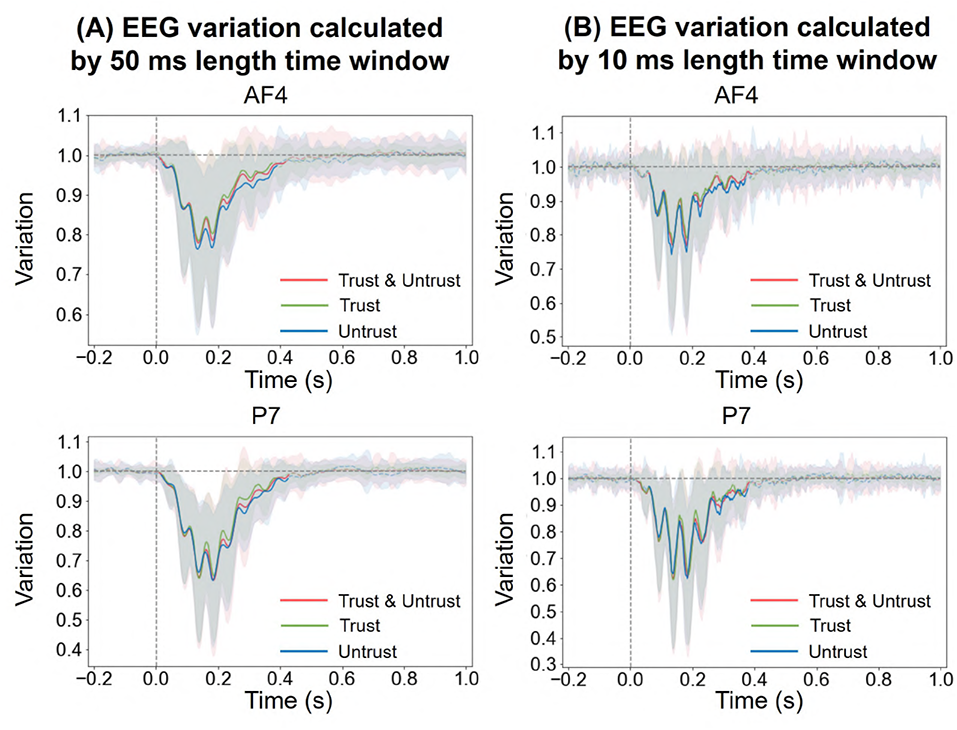
Some representative epochs of the EEG temporal variations calculated by other time windows of different length. The electrode name from upper to bottom: AF4, P7. (A) EEG variations calculated by 50 ms length time window. (B) EEG variations calculated by 10 ms length time window. The red, green and blue line are the variability calculated by all events (Trust & Untrust), only Trust event and only Untrust event, respectively. The shaded areas represent the standard deviation. Dashed lines indicate that the variability is not significantly different from the baseline, and solid lines indicate that the variability is significantly different from the baseline.

**Figure S11:**
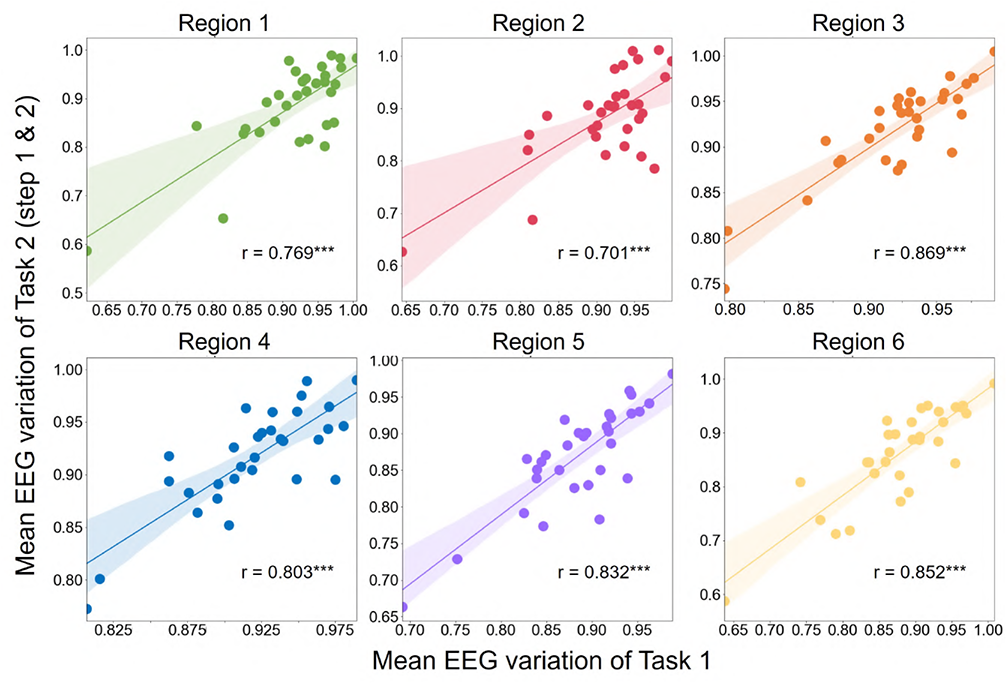
The correlation between the mean EEG variability of Experiment 1 and the mean EEG variability of Experiment 2 & 3. Significance levels are indicated by asterisks: *p *≤* 0.05; **p *≤* 0.01; ***p *≤* 0.001.

